# Genetic modification strategies for electroporation and CRISPR-Cas-based technologies in the non-competent Gram-negative bacterium *Acinetobacter* sp. Tol 5

**DOI:** 10.1101/2023.12.20.572688

**Authors:** Masahito Ishikawa, Katsutoshi Hori

**Author notes:** Corresponding author E-mail address (M. I.) (K. H.).

## Abstract

Environmental isolates are promising candidates for new chassis of synthetic biology because of their inherent conversion capabilities and resilience to environmental stresses; however, many remain genetically intractable and unamenable to established genetic tools tailored for model bacteria. *Acinetobacter* sp. Tol 5 possesses intriguing properties for use in synthetic biology applications. However, genetic manipulation via electroporation is hindered by its low transformation efficiency. This study demonstrated the genetic refinement of the Tol 5 strain, achieving efficient transformation via electroporation. We deleted two genes encoding restriction enzymes. The resulting mutant strain not only exhibited marked efficiency of electrotransformation but also proved receptive to both *in vitro* and *in vivo* DNA assembly technologies, thereby facilitating the construction of recombinant DNA. In addition, we successfully adapted a CRISPR-Cas9-based base-editing platform developed for other *Acinetobacter* species. Our genetic modification strategy allows for the domestication of non-model bacteria, streamlining their utilization in synthetic biology applications.

## Introduction

Synthetic biology designs biological systems for applications beyond the realm of natural evolution. Although *Escherichia coli* has been used as the most tractable chassis over the past two decades, synthetic biology has recently sought a more robust, efficient, and diverse chassis capable of thriving in diverse environments and churning out complexes that *E. coli* can struggle to produce^1, 2^. Genetic manipulation is essential not only for elucidating the biological functions of bacterial genes, but also for effectively employing bacteria in synthetic biology applications encompassing industrial, environmental, and biomedical fields. CRISPR-Cas-based technologies have remarkably enhanced the efficiency of correlating genotype and phenotype^3–5^, and constructed advantageous bacteria for synthetic biology applications^6–9^. However, these techniques exhibit efficient performance in *E. coli* and some closely related enterobacteria, while functioning inefficiently in other bacterial species. Therefore, researchers need to develop a new tool or modify an existing tool for applying CRISPR-Cas-based technologies to bacterial species of interest. Species specificity of CRISPR-Cas-based systems is more pronounced in prokaryotes than in eukaryotes^10^. Transformation and recombination efficiencies with exogenous DNA are postulated factors underlying species specificity. Each bacterial species exhibits a distinct resistance to exogenous DNA. For instance, *Acinetobacter baylyi* ADP1 exhibits high competency, facilitating natural transformation and a strong inclination that is sufficient for homology-directed recombination without the assistance of a phage-derived recombinase.^11^ Conversely, other *Acinetobacter* species do not possess such traits. Non-naturally competent *Acinetobacter baumannii* strains are transformed via electroporation, and their CRISPR-Cas-based genome editing necessitates the use of recombinases derived from phages infecting *Acinetobacter* species^12^. Even within the same *Acinetobacter* species, genetic manipulation and genome editing require different strategies.

*Acinetobacter* sp. Tol 5 is an environmentally isolated Gram-negative bacterium with intriguing characteristics for industrial and environmental applications. Tol 5 can metabolize various chemicals^13, 14^ and exhibits high adhesion to various abiotic surfaces, ranging from hydrophobic plastics to hydrophilic glass and stainless steel through the bacterionanofiber protein AtaA^15^. AtaA-mediated adhesiveness enables Tol 5 cells to function as immobilized biocatalysts for pollutant removal and chemical production^16, 17^. We developed genetic manipulation tools for engineering Tol 5, including a replicable plasmid for expressing genes of interest and suicide plasmids for target gene deletion from Tol 5’s genome^15, 18^. Nevertheless, these tools were designed based on transformation through bacterial conjugation because of the low transformation efficiency of Tol 5 via electroporation. Although bacterial conjugation is commonly employed for the efficient transformation of non-competent bacteria, it requires multiple steps to obtain the desired transformant and gene knockout mutant, rendering genetic manipulations laborious. Furthermore, conjugative plasmids must incorporate the sequence of the origin of transfer. Despite the availability of valuable plasmids from Addgene (https://www.addgene.org/), those devised for competent *Acinetobacter* strains are incompatible with direct engineering of Tol 5 cells. In contrast, electroporation facilitates transformant acquisition and genome editing in fewer steps than those required for bacterial conjugation. Electroporation-based transformation and genome editing would expedite the construction of beneficial Tol 5 strains for synthetic biology applications.

In this study, we reported on a genetically engineered Tol 5 strain capable of efficient transformation through electroporation. Restriction-modification (R-M) systems are recognized as prevalent barriers against foreign DNA invasion by bacteria. Consequently, we hypothesized that the R-M systems in Tol 5 hinder electrotransformation. We identified four putative R-M systems within the Tol 5 genome by sequence similarity searches and excised their restriction enzymes (REases) using suicide plasmids through a conjugation-based approach. The resultant mutant strain not only underwent efficient transformation by electroporation, but also proved amenable to *in vitro* and *in vivo* DNA assembly technologies. Genome editing tools developed for other *Acinetobacter* strains were used to inactivate the gene of Tol 5.

## Results

### Influence of base modifications in a plasmid on electroporation-based transformation

Restriction-modification (R-M) systems play a protective role in bacterial cells by distinguishing between their own DNA and foreign DNA, such as phage DNA, and frequently impede bacterial transformation using recombinant plasmid^19–21^. We previously reported the complete genome sequence of Tol 5, which encompasses 4,402 protein-encoding genes^22^. Sequence similarity search of Tol 5’ genome using REBASE^23^ database (http://rebase.neb.com/rebase/rebase.html), the restriction enzyme database, revealed the presence of four putative R-M systems, including type I, IIG, III, and IV (Table 1). Type I, IIG, and III R-M systems shield host bacterial cells from foreign DNA lacking host-specific base modifications. Conversely, type IV R-M system defends against foreign DNA with specific base modifications. Therefore, we believe that these R-M systems limit the electrotransformation of Tol 5. To validate this hypothesis, electroporation was conducted using an *E. coli*-*Acinetobacter* shuttle plasmid, pARP3, extracted from *E. coli* DH10B or Tol 5 transformant cells (Figure 1). Electroporation with pARP3 sourced from *E. coli* DH10B yielded a transformation efficiency of a mere 3.7 colony-forming units (CFU)/µg-DNA. However, utilizing the pARP3 plasmid sourced from the Tol 5 transformant amplified this value to an impressive 4.1×10^8^ CFU/µg-DNA. *E. coli* DH10B has two DNA methyltransferases that target the N6 position of adenine (m6A): DAM methyltransferase, which methylates adenine in the GA*TC motif, and EcoKI methyltransferase, which methylates the second adenine in the AA*C(N6)GTGC motif and the third adenine in the GCA*C(N6)GTT motif. Yet, Tol 5 cells barely received the pARP3 plasmid replicated in *E. coli* DH10B. The resultant transformation efficiency, which relied on the source of the pARP3 plasmid, implied that the R-M systems in Tol 5 employed adenine modifications divergent from *E. coli* DH10B. Single-molecule real-time (SMRT) sequencing, renowned for its precision in detecting m6A, revealed that the pARP3 plasmid from the Tol 5 transformant contained 26 methylated adenines, some of which deviated from the recognition sequences affiliated with DAM and EcoKI methyltransferase (Table S1). Given that *E. coli* DH10B is deficient in the DCM methyltransferase, the type IV R-M system in the Tol 5 strain is unlikely to target the pARP3 plasmid possessing a C5-methylated cytosine (m5C) within the CCWGG motif. Parallel electroporation experiments using the pARP3 plasmid replicated in *E. coli* DH5α, which harbors both DAM and DCM methylases, mirrored the transformation efficiencies observed with the plasmid from *E. coli* DH1B (Data not shown). Therefore, types I, II, and III R-M systems in the Tol 5 strain may not recognize DNAs possessing m5C within the CCWGG motif. Although SMRT sequencing is less sensitive towards methylated cytosine (m4C) and performs poorly for m5C methylation^24^, we also performed basecalling toward m4C and detected 305 candidates (Table S2). The results of electroporation and SMRT sequencing suggest that unique R-M systems are key limiting factors for the electrotransformation of Tol 5.

**Figure 1.**
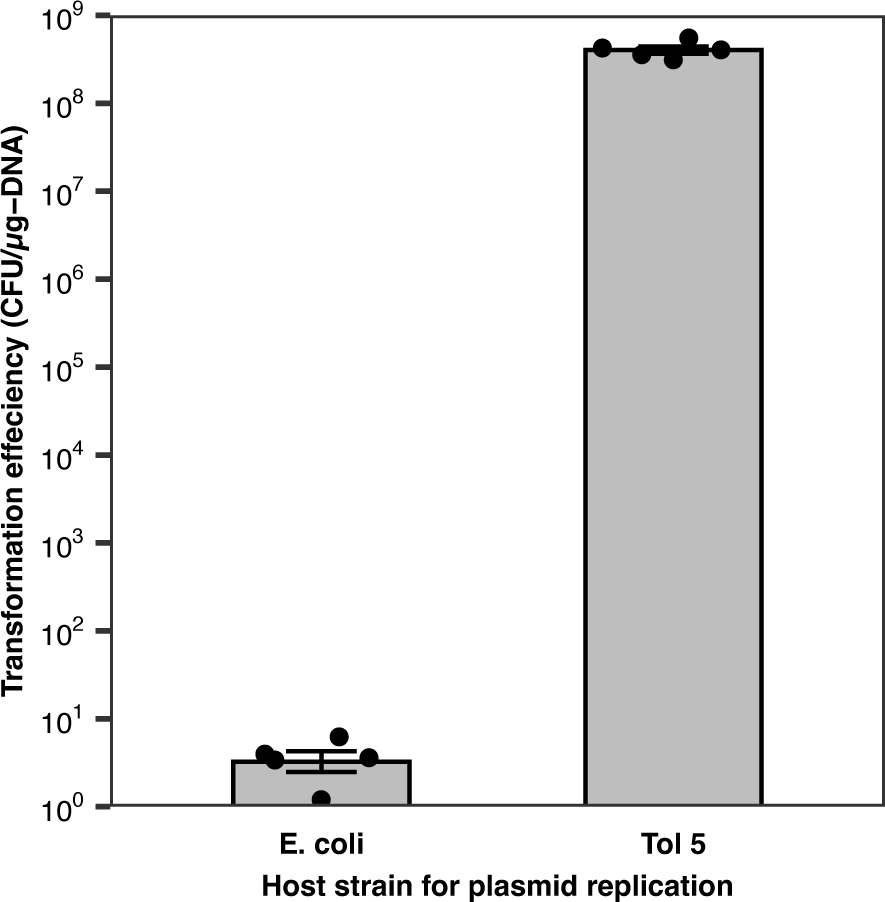
Transformation efficiency of *Acinetobacter* sp. Tol 5 when electroporated with the *E. coli-Acinetobacter* shuttle plasmid pARP3. The pARP3 plasmids, extracted from either *E. coli* DH10B transformants or Tol 5 transformants, were used to electroporate the wild-type Tol 5 strain. Data represent the means from five replicates ± standard deviations.

**Table 1.** Restriction enzymes found in the genome of *Acinetobacter* sp. Tol 5.

| Type | Name | Gene locus tag<sup>a</sup> | Best hit in REBASE<sup>b</sup> |
| --- | --- | --- | --- |
| I | AspTol5ORF4019P | TOL5_40190 | DvuI (AAS96180) |
| IIG | AspTol5ORF3984P | TOL5_39840 | Eco57I (AAA23389) |
| III | AspTol5ORF227P | TOL5_02270 | Vsp69I (CCF17694) |
| IV | AspTol5ORF4146P | TOL5_41460 | ZmoCP4Mrr (ABY40736) |

<sup>a</sup>The genome sequence of *Acinetobacter* sp. Tol 5 was deposited in the DDBJ/GenBank under the accession numbers AP024708 and AP024709.
<sup>b</sup>The restriction enzyme database (http://rebase.neb.com/rebase/rebase.html).

### Identification of restriction nucleases hampering electroporation-based transformation of *Acinetobacter* sp. Tol 5

To investigate the influence of the R-M systems in Tol 5 on the transformation efficiency by electroporation, we constructed mutants deficient in a gene encoding a putative restriction nuclease (REase). Electroporation of these mutants with pARP3 sourced from *E. coli* DH10B yielded an increase in transformation efficiency to 1.0±0.4×10^3^ CFU/µg-DNA and 1.4±0.5×10^2^ CFU/µg-DNA for mutants deficient in type I (REK1) and type III (REK3) REases, respectively (Figure 2). Mutants lacking type II and IV REases (REK2 and REK4) exhibited unaltered transformation efficiency. This observation distinctly implies the role of type I and III R-M systems in the impeding electrotransformation using the *E. coli-*sourced pARP3 plasmid. In contrast, the deletion of type IIG and VI REase genes did not have a similar effect, suggesting the absence of sequence motifs recognized by these REases on the pARP3 plasmid. A combined deficiency in type I and III REases (double mutant, REK13) culminated in a transformation efficiency of 2.1±1.5×10^5^ CFU/µg-DNA, surpassing individual mutant efficiencies. The transformation efficiency of the double mutant REK13 with the pARP3 plasmid replicated in *E. coli* DH10B was approximately 2,000-fold lower than that of the wild-type strain with the pARP3 plasmid replicated in Tol 5 (upper dashed line in Figure 2). This suggests the involvement of methylation-directed defense systems against foreign DNA, distinct from conventional R-M systems in Tol 5. The bacteriophage exclusion (BREX) system, a relatively recently identified defense mechanism^25^, could be an influencing factor. This system, which modifies phage DNA to differentiate it from bacterial DNA, might also target plasmid DNA, as suggested by transformation experiments with the BREX-lacking *Bacillus subtilis* BST7003.^25^ Two sets of gene clusters homologous to components of the BREX system (BCX74278–BCX74285 and BCX76169–BCX76174) were found in the complete genome sequence of Tol 5. The BREX systems in Tol 5 may hamper transformation via electroporation.

**Figure 2.**
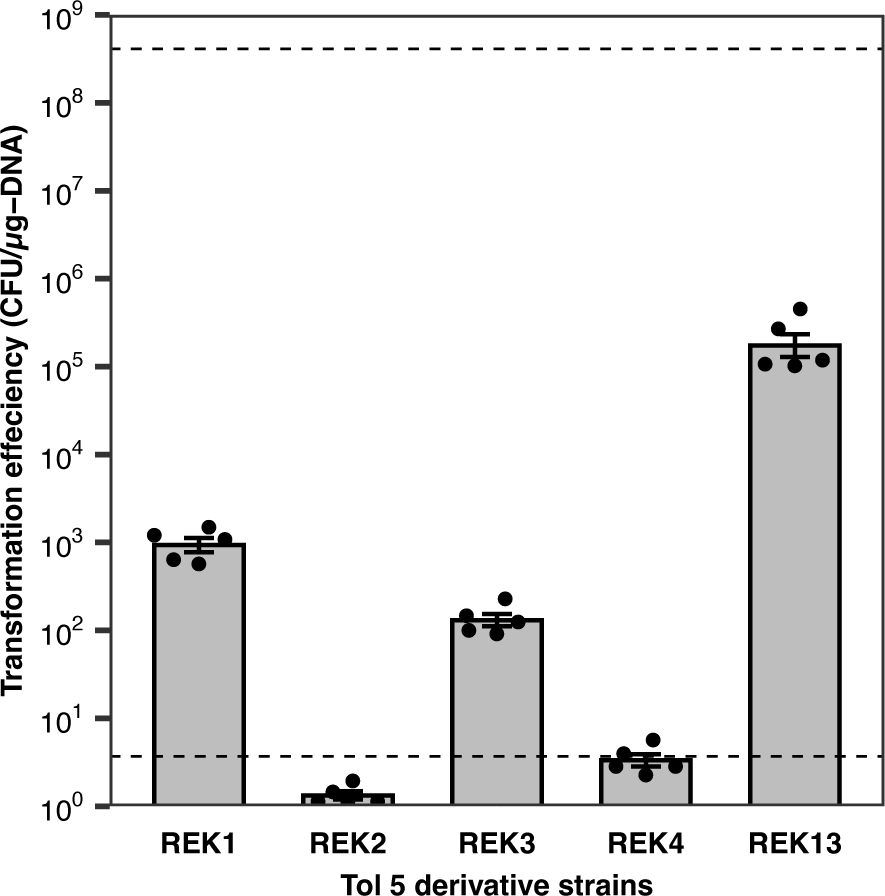
Effect of deleting restriction nuclease genes on the transformation efficiency of *Acinetobacter* sp. Tol 5. The pARP3 plasmid replicated in *E. coli* DH10B was electroporated into Tol 5 mutants lacking genes encoding certain restriction enzymes. Upper dash line represents the transformation efficiency of Tol 5 wild-type strain with pARP3 plasmid replicated in Tol 5, which is shown as the rightmost bar in Figure 1. Lower dash line represents the transformation efficiency of Tol 5 wild-type strain with the pARP3 plasmid replicated in *E. coli* DH10B, the leftmost bar in Figure 1. Data represent the means from five replicates ± standard deviations.

REases play a protective role in bacterial cells by degrading invasive DNAs from phages, transferable plasmids, and transposons. As bacterial cells cannot differentiate recombinant DNAs from these invasive elements, REases eliminate them based on different base modifications. For synthetic biology applications, recombinant DNAs are commonly propagated *in vitro* or within *E. coli* cells. Cell-free DNA amplification technologies, such as PCR and rolling circle amplification (RCA), yield unmethylated products, whereas *E. coli* specifically modifies recombinant plasmid DNA using its own methyltransferases. This presents a challenge when the R-M systems of the target bacteria differ from *E. coli*, complicating the transformation process with recombinant DNAs prepared *in vitro* or within *E. coli* cells. The REBASE database documents various methylation patterns across bacterial species, and metagenomic analysis unveils new methylation motifs, underscoring the diversity of R-M systems.^26^ This diversity explains why bacteria isolated from the environment are often resistant to transformation with recombinant DNAs prepared *in vitro* or within *E. coli* cells.

### *In vitro* and *in vivo* recombinant DNA assembly for *Acinetobacter* sp. Tol 5

For synthetic biology applications in non-model bacteria, recombinant DNA is typically constructed within competent *E. coli* cells using a combination of linearized plasmids and insert DNAs. Subsequently, non-model bacteria are transformed with the resulting recombinant DNA extracted from *E. coli* cells. Constructing recombinant DNA directly within non-model bacteria can expedite the cell engineering process for synthetic biology applications. Commercially available *E. coli* competent cells show the extremely high transformation efficiency, approximately 10^8^–10^10^ CFU/µg-DNA. Although such competent cells are powerful tools for constructing DNA libraries and assembling multiple DNA fragments, routine DNA cloning and assembly do not require extremely high transformation efficiencies. Therefore, the REK13 mutant appears available for constructing recombinant DNA for cell engineering. Modern DNA assembly techniques such as in-fusion cloning, Gibson and Golden Gate assembly have largely superseded traditional DNA cloning, which relies on restriction enzymes and ligases. NEBuilder HiFi DNA assembly, an advancement from Gibson assembly, assembles linear DNA fragments *in vitro* using thermostable DNA polymerase, thermostable DNA ligase, and thermolabile T5 5′→3′ exonuclease. We attempted to transform the Tol 5 REK13 mutant with recombinant DNA assembled *in vitro* using the NEBuilder HiFi DNA assembly (Figure 3A). The vector DNA fragment was amplified by inverse PCR from the pAPR3 plasmid, whereas the insert DNA fragment containing *gfp* gene was amplified by PCR using primers that included 20 bp overlaps of the ends of the vector DNA fragment. After incubation for *in vitro* DNA assembly, the reaction mixture was directly used for electroporation of Tol 5 (WT) or the Tol 5 REK13 mutant (REK13). While the electroporation of WT yielded no colony, the electroporation of the REK13 mutant generated 7.7±0.58×10^5^ colonies per 1 pmol-vector DNA generated (Figure 3B). An impressive 96.7±0.74% of the colonies from the REK13 transformant displayed green fluorescence under blue LED light (Figure 3C), indicating successful DNA assembly. Hence, the Tol 5 REK13 mutant could be effectively transformed with synthetic recombinant DNA assembled *in vitro*, enabling the selection of desired clones from its transformants.

**Figure 3.**
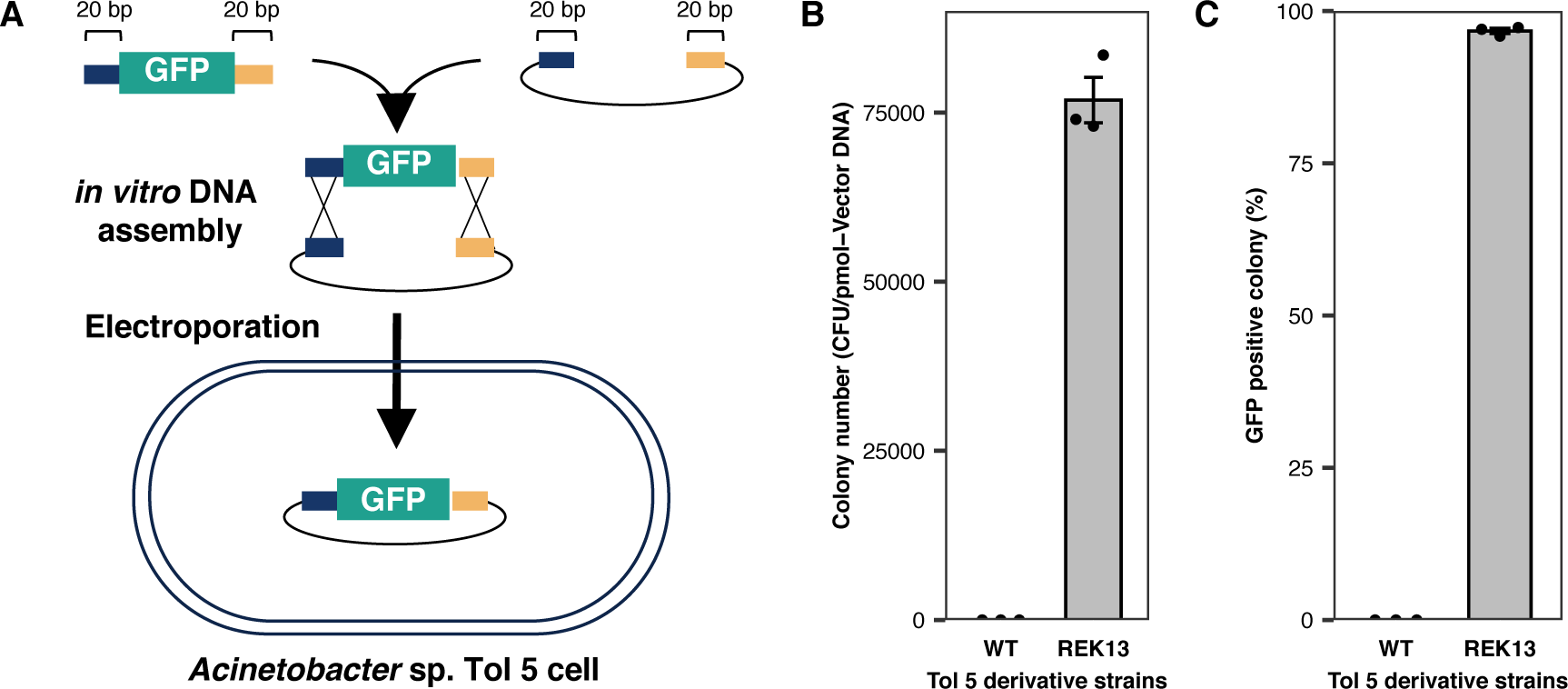
Electrotransformation of *Acinetobacter* sp. Tol 5 cells with an *in vitro* assembled plasmid DNA. (A) Scheme of the electrotransformation of Tol 5 with an *in vitro* assembled DNA. PCR fragments including 20 bp overlaps at their ends undergo *in vitro* using NEBuilder HiFi DNA assembly. The vector PCR fragment carries genes for gentamicin resistance (Gm^R^) and ampicillin resistance (Amp^R^), while the insert PCR fragment includes the *gfp* gene (GFP). The assembly reaction mixture is directly employed for Tol 5 electroporation. (B) Colony number appeared on LB agar plates with gentamycin and ampicillin after the electrotransformation. Data represent the means from three replicates ± standard deviations. (C) Percentage of GFP positive colony among those that appeared on LB agar plates with ampicillin and gentamicin. Data represent the means from three replicates ± standard deviations.

In addition to *in vitro* DNA assembly techniques, *in vivo* DNA assembly techniques have also been used to construct recombinant DNA for synthetic biology applications. The *in vivo E. coli* cloning method (also referred to as iVEC), for instance, provides a more streamlined approach, eschewing the need for costly enzyme solutions.^27, 28^ In typical iVEC methods, linear DNA fragments bearing 20–50-bp overlapping sequences at their ends are simply introduced into *E. coli* cells through calcium chloride treatment followed by heat shock or electroporation. We adapted this *in vivo* cloning method for the Tol 5 REK13 mutant using the iVEC protocol (Figure 4A). Electroporation of DNA fragments with 20 bp overlapping sequences, identical to those used in the NEBuilder HiFi DNA assembly (Figure 3), yielded approximately 1.5±0.58×10^3^ colonies per pmol of vector DNA on Luria-Bertani (LB) agar plates containing ampicillin and gentamycin (Figure 4B). Although the colonies generated by *in vivo* DNA assembly were approximately 51-fold fewer than those generated by *in vitro* assembly, they all exhibited green fluorescence, indicating successful DNA assembly (Figure 4C). Electroporation experiments using DNA fragments with 10 bp and 30 bp overlaps were also conducted. Prolonging overlap sequences to 30 bp augmented the colony count to 4.0±0.98×10^3^ CFU/pmol of vector DNA, with a 98.3±1.6% GFP positivity. Conversely, reducing the overlaps to 10 bp significantly diminished both the colony number (50±86.7 CFU/pmol of vector DNA) and GFP fluorescence positivity (11.1±19.2%). In summary, employing both *in vitro* and *in vivo* DNA assembly techniques with the Tol 5 REK13 mutant eliminated the need for intermediate *E. coli* constructs, thus facilitating the development of tailor-made strains for synthetic biology applications.

**Figure 4.**
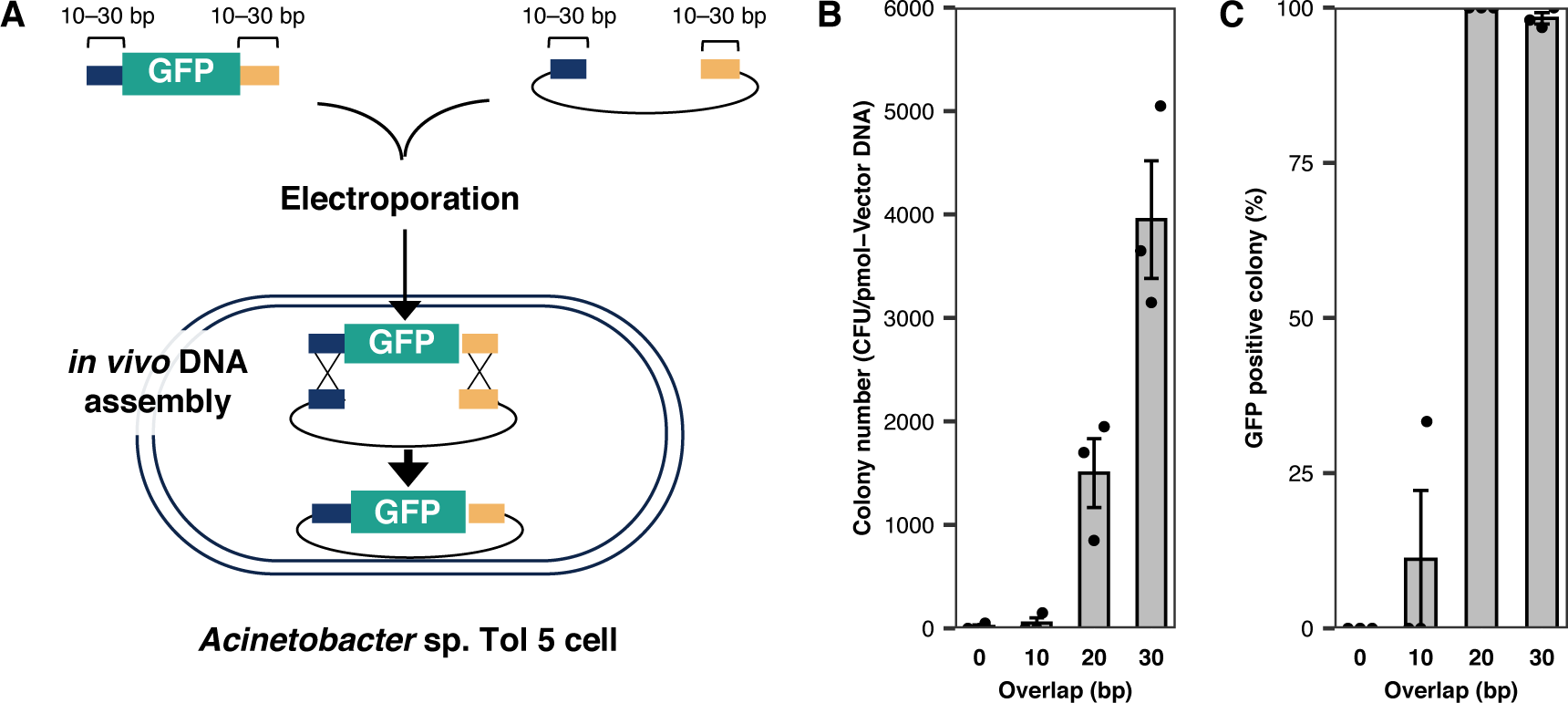
Electroporation-based *in vivo* DNA assembly of *Acinetobacter* sp. Tol 5 strains. (A) Scheme of in vivo DNA assembly of *Acinetobacter* sp. Tol 5. PCR fragments including 10–30 bp overlaps at their ends are electroporated into a Tol 5 mutant strain deficient in type I and III restriction nucleases (REK13). The PCR fragment for an insert contains *gfp* gene (GFP), whereas the PCR fragment for a vector contains gentamycin resistance (Gm^R^) and ampicillin resistance gene (Amp^R^). (B) Colony number appeared on LB agar plates with gentamycin and ampicillin after the electrotransformation. Data represent the means from three replicates ± standard deviations. (C) Percentage of GFP positive colony among those that appeared on LB agar plates with ampicillin and gentamicin. Data represent the means from three replicates ± standard deviations.

### CRISPR-Cas9-based technologies applied to *Acinetobacter* sp. Tol 5

Genome engineering, in addition to plasmid DNA transformation, is crucial for constructing bacteria that are useful for synthetic biology applications. Various CRISPR-Cas9 based technologies with species-specific modifications have been developed for deleting and inserting genes from diverse bacteria.^29^ Wang et al. developed highly efficient CRISPR-Cas9 based genome and base-editing methods optimized for *Acinetobacter baumannii.*^12^ Although the CRISPR-Cas9 based systems for *A. baumannii* were likely to function in Tol 5, implementation was deterred because of the requirement for electrotransformation with plasmids carrying Cas9 and sgRNA genes. However, the deletion of the two restriction enzyme genes facilitated the transformation of the Tol 5 REK13 mutant with plasmid DNA through electroporation. Thus, we tested whether the CRISPR-Cas9 system for *A. baumannii* could cleave the genome of the Tol 5 REK13 mutant. Tol 5 REK13 mutant cells harboring the pCasAb-apr plasmid (containing Cas9 nuclease) were transformed with the pSGAb-km_*ataA* plasmid containing the 20-bp spacer sequence of *ataA* or empty pSGAb-km. Electroporation with the empty pSGAb-km yielded 4426±2753 colonies, whereas that with pSGAb-km_*ataA* yielded a mere 9.5±8.2 colonies (Figure 5). This result implies that the Cas9-sgRNA complex targets and cleaves to the Tol 5 genome, and that the subsequent double-strand break (DSB) is lethal. Subsequently, we electroporated pSGAb-km_*ataA*_HR, containing the 20-bp spacer sequence of *ataA* and the repair template of *ataA* lacking a 274-bp sequence for homologous recombination, into Tol 5 REK13 mutant cells harboring the pCas9-apr plasmid. However, no difference was observed between electrotransformation with and without the repair template, suggesting that there was no homologous recombination after the Cas9-induced DSB. Colony PCR revealed no 274-bp deletion through homologous recombination with the repair template after Cas9-induced DSB. Although genome editing was not performed, the lethality of the Cas9-induced DSB confirmed the binding and cleaving activity of the Cas9-sgRNA complex on the genomic DNA of Tol 5.

**Figure 5.**
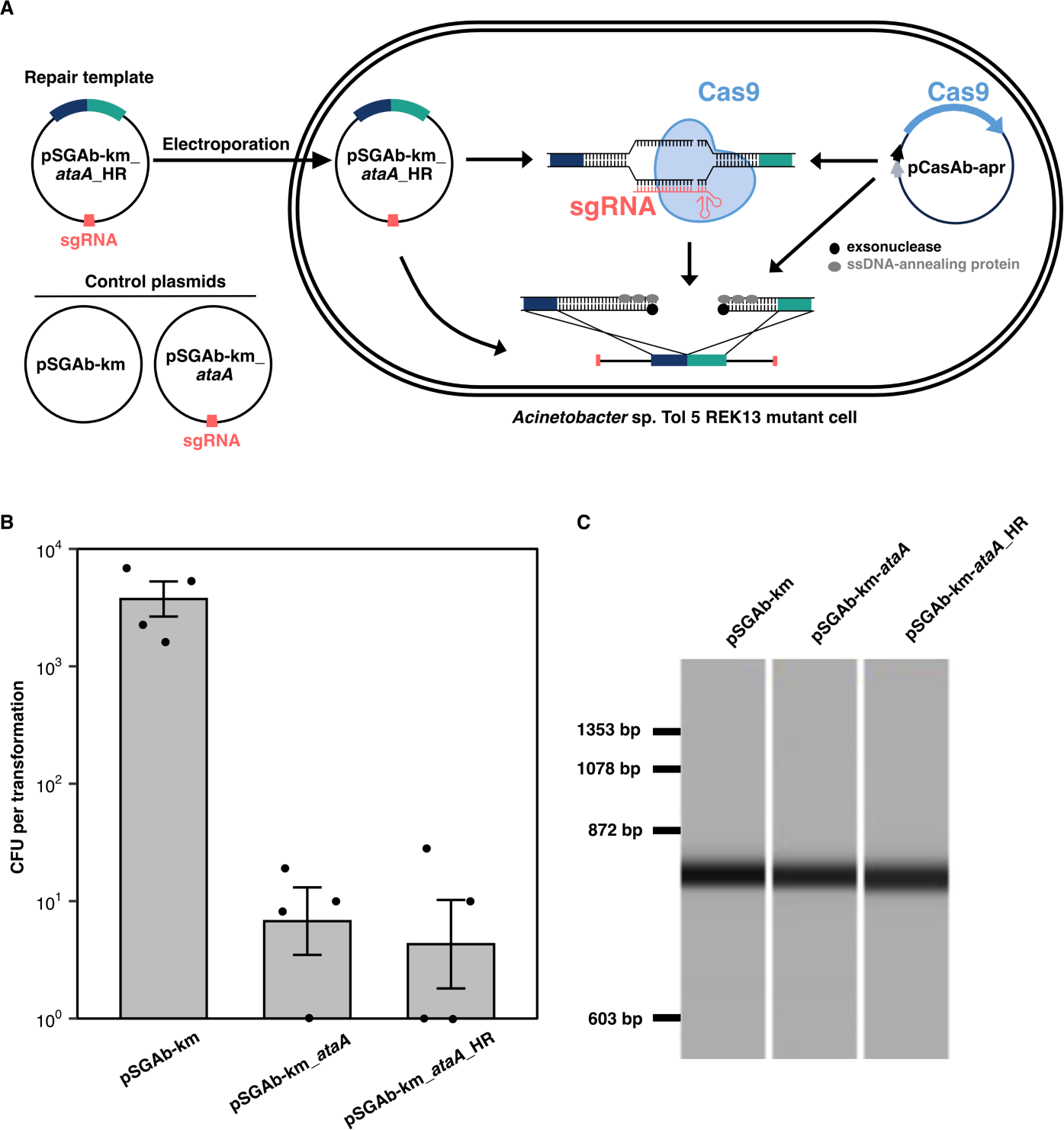
CRISPR-Cas9-based genome editing in *Acinetobacter* sp. Tol 5. (A) Scheme of CRISPR-cas9-based genome editing in *Acinetobacter* sp. Tol 5 REK13 mutant. The pSGAb-km_*ataA*_HR plasmid contains the sgRNA that directs Cas9 to the *ataA* gene and a repair template for homologous recombination following a Cas9-sgRNA-induced double-strand break (DSB). The pCasAb-apr plasmid supplies Cas9, which creates DSBs in the Tol 5 REK13 mutant’s genome in concert with sgRNA. The pSGAb-km_*ataA*_HR plasmid is also cleaved by the Cas9-sgRNA complex and subsequently serves as a template for homologues recombination to repair the genome injured by the DSB. (B) The colony-forming unit (CFUs) of each transformation using different plasmids. The pSGAb-km plasmid, which harbors none of the sgRNA nor repair template, was used as a negative control for the genome editing of *ataA*. (C) Representative gel-like images of colony PCR amplicons displayed on the MCE-202 MultiNA. The attempted deletion of the *ataA* gene was examined using the colony PCR using primers annealing to the outside of the flanking regions of the DSB used as homologous sites. No successful deletion was confirmed in the twelve tested colonies.

Wang et al. also developed a CRISPR-Cas9-based base-editing system for *A. baumannii*^12^. The utilization of APOBEC1, which is merged with Cas9 (D10A) nickase, is unique to this system and allows for the conversion of cytidine to thymidine without introducing a DSB. This bypasses the need for homologous recombination for targeting specific genes. We applied this CRISPR-Cas9-based base-editing to introduce a stop codon within *ataA* in the Tol 5 REK13 mutant (Figure 6A). Notably, the 20 nt-spacer sequence of *ataA* remained consistent with that used for genome editing (Figure 6B). Electroporation of the Tol 5 REK13 mutant was conducted with pBECAb-apr_*ataA* (consisting of APOBEC1 fused to Cas9 (D10A) nickase and sgRNA), yielding several hundred colonies on an LB agar plate supplemented with apramycin. Subsequent colony PCR and sequencing identified specific colonies carrying the desired mutations that inhibited *ataA* transcription. The resultant DNA sequencing results revealed that the glutamic acid codon (CAA) at the 116th amino acid position in AtaA was switched to the stop codon (TAA) (Figure 6C).

**Figure 6.**
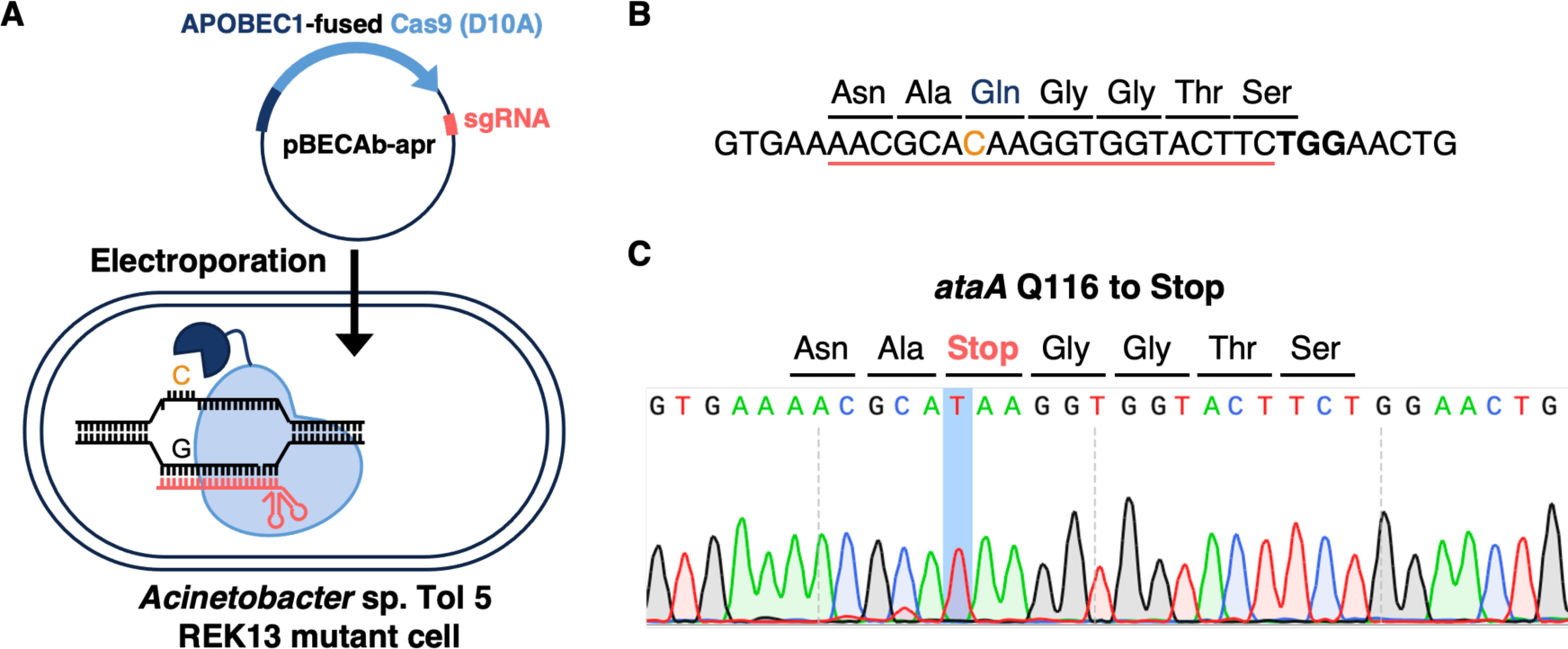
CRISPR-Cas9-based base editing. (A) Scheme of CRISPR-Cas9-based base editing in *Acinetobacter* sp. Tol 5 REK13 mutant. The pBECAb-apr plasmid supplies the APOBEC1-fused Cas9 (D10A) nickase in Tol 5 REK13 mutant cells. The APOBEC1-fused Cas9 editor converts C to U, finally forming the T:A base pair. (B) Design of the spacer sequence targeting the *ataA* gene for knockout. The spacer sequence intended for a stop codon insertion is highlighted by red underline. The targeted cytosine within the glutamic acid codon (Q116) is indicated in orange. The PAM sequence is highlighted in bold. (C) A representative sequencing result from a base-edited Tol 5 REK13 mutant colony.

The base-editing method successfully inactivated the *ataA* gene; however, attempts at genome editing to remove it were unsuccessful. This may be due to the incompatibility of the recombination system within the pCasAb-apr plasmid with Tol 5 cells. When Wang et al. developed the pCasAb-apr plasmid, they tested three different exogenous recombination systems (Red from lambda phage, RecET from Rac prophage, and RecAb from *A. baumannii*) and identified that only RecAb could repair the DSBs generated by CRISPR-Cas9 in *A. baumannii.*^12^ For successful genome editing, we need to find an exogenous recombination system that facilitates homologous recombination in Tol 5 cells. As many bacteria lack the ability to perform non-homologous end-joining, species specificity of the recombination system is a critical consideration when applying CRISPR-Cas9-based technologies to non-model organisms.

### Conclusion

In this study, by deleting two genes encoding REases, which form part of the typical bacterial defense systems against foreign DNA, we successfully introduced recombinant DNA into Tol 5 cells with high efficiency. This advancement facilitated the use of *in vivo/in vitro* DNA assembly technologies and the CRISPR-Cas-based base-editing method previously developed for different *Acinetobacter* strains. The ease of genetic manipulation in laboratory *E. coli* strains has significantly propelled synthetic biology research over the past two decades. However, these strains do not always serve as ideal hosts for industrial and environmental applications, prompting the search for new bacterial chassis.^1, 2^ Although new chassis for synthetic biology need to be genetically tractable, many environmental isolates remain intractable. Phages are the most abundant biological entities on Earth, with the particle count estimated to exceed 10 times that of bacteria.^30^ Given the prevalence of phages, environmental bacteria have evolved robust defense systems against them. These defense systems often target and eliminate recombinant DNAs constructed *in vitro* or within *E. coli* cells. Methods for *in vivo/in vitro* DNA assembly directly using target bacteria eliminate the need for intermediate *E. coli* constructs. Our approach to the genetic modification of non-competent bacteria paves the way for the domestication of beneficial non-model bacteria, advancing their application in synthetic biology.

## Methods

### Bacterial culture

*Acinetobacter* sp. Tol 5 and its derivative strains were grown in basal salt (BS) medium supplemented with 0.05% (vol/vol) toluene or LB medium as previously described.^15^ *Escherichia coli* strains were grown in LB medium containing the appropriate antibiotics at 37°C. Antibiotics were used at the following concentrations, when required: ampicillin (100 μg/ml), apramycin (100 µg/mL), gentamicin (10 μg/ml), and kanamycin (50 μg/ml).

### Construction of a deletion mutant in *Acinetobacter* sp. Tol 5

Deletion mutants of Tol 5 presented in Table S3 were constructed using the plasmid-based counterselection method as described previously.^31^ The primers and plasmids used for the construction of mutant strains are presented in Tables S4 and S5, respectively. Briefly, approximately1-kb regions upstream and downstream regions of a target gene were amplified by PCR and assembled with BamHI-cut pJQ200sk using the NEBuilder HiFi DNA assembly system (New England Biolabs, Ipswich, MA) to generate a suicide plasmid. Tol 5 or its derivative mutant was mated with *E. coli* S17-1 harboring the suicide plasmid on an LB agar plate for 24 h at 28°C. The cells were collected in 1 mL of 0.85% NaCl solution, plated on BS agar plates containing gentamicin (100 µg/mL), and incubated with toluene vapor for 2 days at 28°C. Chromosomal integration of the suicide plasmid was confirmed by colony PCR using primers annealing to the outside and inside of the flanking region of the target gene, which were used as homologous sites for recombination. For the deletion of a target gene, the resulting single-crossover mutant was spread on a BS agar plate containing 5% sucrose and incubated with toluene vapor for 2 days at 28°C. Deletion of the target gene was confirmed by PCR using primers annealing to the outside of the flanking region of the target gene used as homologous sites for recombination.

### Electroporation

Tol 5 and its derivative mutants were inoculated into 5 mL of 2×YT medium in a 50-mL conical tube and incubated at 28°C with shaking at 125 rpm. After overnight incubation, the cells were harvested by centrifugation at 8,000×g for 3 min at 4°C, rinsed with 10 mL 10% (wt/v) glycerol solution three times, and resuspended in 500 µl 10% (wt/v) glycerol solution. One hundred microliters of the cell suspension was mixed with DNA and transferred to an ice-cold 2-mm-gap cuvette. Electroporation was performed using a Gene Pulser II system (Bio-Rad, Hercules, CA) set at 25 µF, 3000 V, and 200 Ω. Immediately after electroporation, 0.9 mL of 2×YT medium at room temperature was added to the cells. The resuspended cells were transferred into a 15-mL conical tube and incubated at 28°C for 2 h with shaking at 125 rpm. After incubation, the cells were spread on a selective plate.

### SMRT sequencing

The pARP3 plasmid for methylation analysis was extracted from 200 mL of an overnight culture of Tol 5 transformant cells using the HiSpeed Plasmid Midi kit (QIAGEN) according to the manufacturer’s instructions. The extracted plasmid was linearized by SalI digestion and purified using the Wizard SV PCR cleanup system (Promega, Madison, WI). SMRT sequencing was performed using Sequel IIe (Pacific Biosciences, Menlo Park, CA). The pARP3 sequence was determined using reads covering the full-length plasmid. Reads were mapped to the pARP3 sequence with pbmm2 1.4.0 in SMRT Tools (Pacific Biosciences). Detection of DNA methylation signatures was performed using ipdSummary in the SMRT Tools program (Pacific Biosciences) with the-- identify m6A, m4C, --methylFraction, and --maxAligments 15000 options.

### In vitro and in vivo DNA assembly

For *in vitro* and *in vivo* DNA assembly, DNA fragments were amplified using KOD-Plus-Ver.2 (Toyobo, Osaka, Japan) and the primers are presented in Table S5. All the resulting amplicons were purified using the Wizard SV PCR cleanup system (Promega). The DNA fragment for the vector was amplified by PCR using the primer set Inv-pARP3-fwd/Inv-pARP3-rev and the pARP3 plasmid as the DNA template. A DNA fragment containing a constitutive promoter, a ribosome-binding site, and *gfp* gene was artificially synthesized as a template for amplifying DNA fragments for inserts.

For *in vitro* assembly, the DNA fragment containing 20-bp-overhangs at the ends was amplified using the primer set 20bp-eGFP-fwd/20bp-eGFP-rev. After purification, 0.18 pmol of the insert DNA fragment and 0.02 pmol of the vector DNA fragment were mixed with NEBuilder HiFi DNA Assembly Master Mix (New England Biolabs) to make a total volume of 10 µl. After 1-h-incubation at 50°C, 1 µl of the reaction mixture was electroporated into Tol 5 or REK13 mutant cells. Transformants were selected on LB agar plates containing ampicillin and gentamicin.

For *in vivo* assembly, the DNA fragments containing 10 bp-, 20 bp-, and 30 bp-overhangs at the ends were amplified using primer sets 10bp-eGFP-fwd/10bp-eGFP-rev, 20bp-eGFP_fwd/20bp-eGFP-rev, and 30bp-eGFP-fwd/30bp-eGFP-rev, respectively. After the purification using Wizard SV PCR cleanup system (Promega), 0.18 pmol of the insert DNA fragment and 0.02 pmol of the vector DNA fragment were directly electroporated into REK13 mutant cells. Transformants were selected on LB agar plates containing ampicillin and gentamicin.

### Genome editing and base editing

CRISPR-Cas9-based genome and base editing were performed according to the methods described by Wang et al^12^. Briefly, two oligonucleotides (5’-TAGTAACGCACAAGGTGGTACTTC-3’ and 5’-TTGCGTGTTCCACCATGAAGCAAA-3’) were synthesized for 20-bp spacer sequence before a protospacer adjacent motif (PAM) site (5’-NGG-3’) in the *ataA* gene. The two oligonucleotides were phosphorylated, annealed, and cloned into the BsaI sites of pSGAb-km and pBECAb-apr using a Golden Gate Assembly Kit (New England Biolabs) to generate pSGAb_*ataA* and pBECAb_*ataA*, respectively. To prepare the plasmid-based repair template of *ataA* for homologous recombination after Cas9-mediated DSB, 214-bp upstream and 284-bp downstream of the DSB site in *ataA* were amplified by PCR using the primer sets RTemp-upst_fwd/RTemp-upst_rev and RTemp-dwst_fwd/RTemp-dwst_rev, respectively. The resulting amplicons were assembled with SalI- and KpnI-cut pSGAb-*ataA* using the NEBuilder HiFi DNA assembly kit (New England Biolabs) to generate pSGAb-*ataA*_HR.

For genome editing, pCasAb-apr was electroporated into Tol 5 REK13 mutant. The resulting transformant was inoculated into 5 mL of 2×YT medium containing 100 µg/mL apramycin in a 50-mL conical tube and incubated at 28°C with shaking at 125 rpm. After overnight cultivation, the cells were prepared for electroporation as described above. Then, 0.1 pmol of pSGAb-km, pSGAb-*ataA*, or pSGAb-*ataA*_HR were electroporated into Tol 5 REK mutant cells harboring pCas9Ab-*apr*. Electroporated cells were plated on LB agar plates containing apramycin, kanamycin, and IPTG (0.1 mM) and incubated at 28°C overnight. Deletion of the *ataA* gene was examined by colony PCR using the primers Edit-Check-F and Edit-Check-2R. Colony PCR amplicons were loaded onto an MCE-202 MultiNA (Shimadzu, Kyoto, Japan).

For base editing, pBECAb-*ataA* was electroporated into the Tol 5 REK13 mutant. Electroporated cells were plated onto LB agar plates containing apramycin. Base editing of the *ataA* gene was confirmed by colony PCR using the primer set Edit-Check-F/Edit-Check-R and subsequent amplicon sequencing. Successfully base-edited mutants were cultivated in LB medium and streaked onto an LB agar plate containing 5% sucrose to cure the pBECAb-*ataA*-spacer plasmid using SacB-based counterselection.

## Acknowledgements

We thank Eriko Kawamoto, Yoko Yamamoto, Umechiyo Matsumura, and Kazuyo Funatsu for technical assistance. This study was supported by PRESTO (grant number JPMJPR20K2) from the Japan Science and Technology Agency.

## Author contributions

M.I. designed the study, performed the experiments, analyzed the data, and wrote the manuscript. K.H. analyzed the data and reviewed the manuscript.

