## Supplemental Table 3-5 for "Genetic modification strategies for electroporation and CRISPR-Cas-based technologies in the non-competent Gram-negative bacterium *Acinetobacter* sp. Tol 5"

*Corresponding author

**Table S3. Bacterial strains used in this study**

| Strain | Description | Reference |
|---|---|---|
| *Acinetobacter* sp. Tol 5 | | |
| WT | Wild-type strain | [1] |
| Δ*ataA* | Tol 5 mutant deficient in the *ataA* gene | [2] |
| REK1 | Tol 5 mutant deficient in the gene encoding the putative type I endonuclease subunit R (AspTol5ORF4019P) | This study |
| REK2 | Tol 5 mutant deficient in the gene encoding the putative type IIG restriction enzyme/methyl transferase (AspTol5ORF3984P) | This study |
| REK3 | Tol 5 mutant deficient in the gene encoding the putative type III endonuclease subunit R (AspTol5ORF227P) | This study |
| REK4 | Tol 5 mutant deficient in the gene encoding the putative type IV methyl-directed restriction enzyme (AspTol5ORF4146P) | This study |
| REK13 | REK1 mutant deficient in the gene encoding AspTol5ORF227P | This study |
| REK13Δ*ataA* | REK13 mutant deficient in the *ataA* gene | This study |
| *Escherichia coli* | | |
| DH5α | Host for routine cloning | Takara Bio |
| DH10B | Host for routine cloning | Invitrogen |
| S17-1 | Donor strain for bacterial conjugation | [3] |

**Table S4. Plasmids in this study**

| Plasmid | Description | Reference |
|---|---|---|
| pJQ200sk | Suicide plasmid, $Gm^R$, SacB | [4] |
| pJQ200RE1KO | A DNA fragment containing the upstream and downstream regions of AspTol5ORF4019P ligated into the BamHI site of pJQ200sk | This study |
| pJQ200RE2KO | A DNA fragment containing the upstream and downstream regions of AspTol5ORF3984P ligated into the BamHI site of pJQ200sk | This study |
| pJQ200RE3KO | A DNA fragment containing the upstream and downstream regions of AspTol5ORF227P ligated into the BamHI site of pJQ200sk | This study |
| pJQ200RE4KO | A DNA fragment containing the upstream and downstream regions of AspTol5ORF4146P ligated into the BamHI site of pJQ200sk | This study |
| pARP3 | *E.coli-Acinetobacter* shuttle plasmid, *araC*-$P_{BAD}$, $Amp^R$, $Gm^R$ | [5] |
| pBECAb-apr | Broad-host-range plasmid harboring the APOBEC1 fused to Cas9(D10A) nickase, $Apr^R$, SacB | [6] |
| pBECAb-apr_*ataA* | pBECAb-apr plasmid harboring the *ataA* spacer sequence | This study |
| pCasAb-apr | Broad-host-range plasmid optimized for the genome editing in *Acinetobacter baumannii*, Cas9, RecAb, $Apr^R$, SacB | [6] |
| pSGAb-km | *E. coli-Acinetobacter* shuttle plasmid, sgRNA, $Km^R$, SacB | [6] |
| pSGAb-km_*ataA* | pSGAb-km plasmid harboring the *ataA* spacer sequence | This study |
| pSGAb-km_*ataA*_HR | pSGAb-km plasmid harboring the *ataA* spacer sequence and the repair template for the homologous recombination of *ataA* | This study |

**Table S5. Primers used for constructing plasmids in this study**

| Primer | Sequence (5'→3') |
|---|---|
| RE1-upst_fwd | CGAATTCCTGCAGCCCGGGGACCAAACGCATTAATGATG |
| RE1-upst_rev | CACTTGAGTCTTAGTCAACAAGGTTAAAGC |
| RE1-dwst_fwd | TGTTGACTAAGACTCAAGTGATGCCTGATTG |
| RE1-dwst_rev | AGAACTAGTGTAGGCTGGAACTTGAATCAC |
| RE2-upst_fwd | CGAATTCCTGCAGCCCGGGGTCCCCTAAAGAGAAGATTTC |
| RE2-upst_rev | CTCGGGTTTACTTAGAGGTACTAAATATCGAG |
| RE2-dwst_fwd | TACCTCTAAGTAAACCCGAGCTTGATGAG |
| RE2-dwst_rev | CGGCCGCTCTAGAACTAGTGAATAATAATGGTGATGCCG |
| RE3-upst_fwd | CGAATTCCTGCAGCCCGGGGGATCATGTCATCAATACTGCGG |
| RE3-upst_rev | GGCAAAACCCAGTATTCGGATTAAACTTCGAC |
| RE3-dwst_fwd | TAATCCGAATACTGGGTTTTGCCTTAGC |
| RE3-dwst_rev | CGGCCGCTCTAGAACTAGTGGATCCTTTGTCCCACCACTATTTG |
| RE4-upst_fwd | CGAATTCCTGCAGCCCGGGGTTTCCGTAGCCGCTTTAAC |
| RE4-upst_rev | CGATTTGATGTCTGCAAAGAAATCCGGAATTG |
| RE4-dwst_fwd | TCTTTGCAGACATCAAATCGTAGTGGGTATAC |
| RE4-dwst_rev | CGGCCGCTCTAGAACTAGTGTTGCAGGTGAAGTTGATC |
| Inv-pARP3-fwd | TTGGGCTAGCGAATTCCTGC |
| Inv-pARP3-rev | TACCCAATTCGCCCTATAGTG |
| 0bp-eGFP-fwd | ACAAGCAACAGTATTCAG |
| 0bp-eGFP-rev | TTATTTATACAGTTCATCCATACC |
| 10bp-eGFP-fwd | GAATTGGGTAACAAGCAACAGTATTCAG |
| 10bp-eGFP-rev | GCTAGCCCAATTATTTATACAGTTCATCCATACC |
| 20bp-eGFP-fwd | ACTATAGGGCGAATTGGGTAACAAGCAACAGTATTCAG |
| 20bp-eGFP-rev | GCAGGAATTCGCTAGCCCAATTATTTATACAGTTCATCCATACC |
| 30bp-eGFP-fwd | AATACGACTCACTATAGGGCGAATTGGGTAACAAGCAACAGTATTCAG |
| 30bp-eGFP-rev | CCCCCGGGCTGCAGGAATTCGCTAGCCCAATTATTTATACAGTTCATCCATACC |
| RTemp-upst_fwd | GACGCGTATTGGGATGGTACATTTGGAATGCGACTTTG |
| RTemp-upst_rev | GGATATTGCTAATTCGTAGAAGCTGTAGC |
| RTemp-dwst_fwd | TCTACGAATTAGCAATATCCAGACAAGC |
| RTemp-dwst_rev | CTGATGCCGTATCGATACCGTTACCGATCCTGCACCTAG |
| RTemp-AtaAF | ATTTGGAATGCGACTTTGTTGG |
| RTemp-AtaAR | TGCCCCATAGCAATCGATCC |
| Edit-Check-F | TTTGTTGGCATGGGTTGCAG |
| Edit-Check-R2 | AGCTGATGCACCTACACCAG |
